# Mixed-polarity random-dot stereograms alter GABA and Glx concentration in the early visual cortex

**DOI:** 10.1101/549980

**Authors:** Reuben Rideaux, Nuno Goncalves, Andrew E Welchman

**Affiliations:** Department of Psychology, University of Cambridge, CB2 3EB, UK

## Abstract

The offset between images projected onto the left and right retinae (binocular disparity) provides a powerful cue to the three-dimensional structure of the environment. It was previously shown that depth judgements are better when images comprise both light and dark features, rather than only dark or only light elements. Since Harris and Parker (1995) discovered the “mixed-polarity benefit”, there has been limited evidence supporting their hypothesis that the benefit is due to separate bright and dark channels. Goncalves and Welchman (2017) observed that single- and mixed-polarity stereograms evoke different levels of positive and negative activity in a deep neural network trained on natural images to make depth judgements, which also showed the mixed-polarity benefit. Motivated by this discovery, here we seek to test the potential for changes in the balance of excitation and inhibition that are produced by viewing these stimuli. In particular, we use magnetic resonance spectroscopy to measure Glx and GABA concentration in the early visual cortex of adult humans while viewing single- and mixed-polarity random-dot stereograms (RDS). We find that observers’ Glx concentration is significantly higher while GABA concentration is significantly lower when viewing mixed-polarity RDS than when viewing single-polarity RDS. These results indicate that excitation and inhibition facilitate processing of single- and mixed-polarity stereograms in the early visual cortex to different extents, consistent with recent theoretical work (Goncalves & Welchman, 2017).

## INTRODUCTION

Binocular stereopsis is one of the primary cues for three-dimensional (3D) vision. It remains an important challenge to understand how the brain combines a pair of two-dimensional retinal images to support 3D perception. A clue to understanding the neural computation of binocular stereopsis may be found in the observation that depth judgements are more accurate when binocular images are composed of both light and dark features, rather than just one or the other (Harris & Parker, 1995).

This ‘mixed-polarity benefit’ was original explained on the basis that bright and dark features are processed by separate ON and OFF channels (Harris & Parker, 1995). Such neural infrastructure would reduce the number of potential binocular matches in a mixed-polarity stimulus, i.e., a random-dot stereogram (RDS), by as much as half, simplifying the stereoscopic correspondence problem considerably. Separate of ON and OFF channels first appear at the bipolar cell level as ON and OFF ganglia (Nelson, Famiglietti, & Kolb, 1978) and are maintained at the retinal ganglion and lateral geniculate nucleus level as ON and OFF centre cells. However, the convergence of ON and OFF channels in V1 to form simple cells (Schiller, 1992) seems to contradict this as a potential explanation for the mixed-polarity benefit.

Recently, Goncalves and Welchman (2017) showed that it is possible to capture the mixed-polarity benefit using a simple linear - nonlinear processing architecture that did not depend on separate ON and OFF channels. Thereafter, Read and Cumming (2018) proposed that the ‘mixed polarity’ benefit could arise from subtle changes in image correlation that can occur in some circumstances. As current models of binocular processing are based on cross-correlation between the left and right eyes (Ohzawa, DeAngelis, & Freeman, 1990), they do not consider image features, but instead compute the inter-ocular cross-correlation between left and right images. Within this framework, higher image correlation might be expected to drive binocular cells in the primary visual cortex more strongly; thus, subtle changes in inter-ocular cross-correlation provide an explanation for the benefit that is consistent with this model. While these theoretical explanations appear to capture the improved behavioural performance associated with mixed-polarity images, empirical evidence is needed to establish the neural basis of the effect.

Here we seek to test the potential for changes in the balance of excitation and inhibition that are produced by viewing these stimuli. This follows directly from observing that single- and mixed-polarity RDS evoke different levels of positive and negative activity in a deep neural network, which also showed the mixed-polarity benefit (Goncalves & Welchman, 2017). In particular, we use magnetic resonance spectroscopy (MRS) to measure primary inhibitory neurotransmitter γ-aminobutyric acid (GABA) concentration in the early visual cortex of human observers while viewing single- and mixed-polarity RDS. We find that viewing single- and mixed-polarity RDS produces differences in the concentration of GABA. Further, we find that that viewing single- and mixed-polarity RDS also produces differences in Glx, i.e., a complex comprising primary excitatory neurotransmitter glutamate and glutamine.

## METHODS

### Participants

Twenty healthy observers from the University of Cambridge with normal or corrected-to-normal vision participated in the MR spectroscopy experiment. The mean age was 25.5 yr (range = 19.4–40.5 yr; 12 women). Participants were screened for stereoacuity (<1° arcmin near/far discrimination threshold) and contraindications to MRI prior to the experiment. All experiments were conducted in accordance with the ethical guidelines of the Declaration of Helsinki and were approved by the University of Cambridge STEM and all participants provided informed consent.

### Apparatus and stimuli

Stimuli were programmed and presented in MATLAB (The MathWorks, Natick, MA) with Psychophysics Toolbox extensions (Brainard, 1997; Pelli, 1997). Stereoscopic presentation in the scanner was achieved using a “PROPixx” DLP LED projector (VPixx Technologies) with a refresh rate of 120 Hz and resolution of 1920 × 1080, operating in RB3D mode. The left and right images were separated by a fast-switching circular polarization modulator in front of the projector lens (DepthQ; Lightspeed Design). The onset of each orthogonal polarization was synchronized with the video refresh, enabling interleaved rates of 60 Hz for each eye’s image. MR-safe circular polarization filter glasses were worn by subjects in the scanner to dissociate the left and right eye’s view of the image. Stimuli were back-projected onto a polarization-preserving screen (Stewart Filmscreen, model 150) inside the bore of the magnet and viewed via a front-surfaced mirror attached to the head coil and angled at 45° above the observers’ heads. This resulted in a viewing distance of 82 cm, from which all stimuli were visible within the binocular field of view.

Stimuli consisted of random-dot stereograms (RDS; 12°×12°) on a midgray background surrounded by a static grid of black and white squares intended to facilitate stable vergence. Dots in the stereogram followed a black or white Gaussian luminance profile, subtending 0.07° at half maximum. There were 108 dots/deg^2^, resulting in ~38% coverage of the background. In the centre of the stereogram, 4 wedges were equally distributed around a circular aperture (1.2°), each subtending 10° in the radial direction and 70° in polar angle, with a 20° gap between wedges (**Fig. 1a**). Dots comprising the wedges were offset by 10 arcmin between the left and right eyes, while the remaining dots had zero offset. Stimuli were presented for 1.8 s and separated by 0.2 s inter-stimulus-intervals consisting of only the background and fixation cross. To reduce adaptation, we applied a random polar rotation on each presentation to the set of wedges such that the disparity edges of the stimuli were in different locations for each stimulus presentation (i.e., a rigid body rotation of the four depth wedges together around the fixation point). Every five presentations we reversed the sign of the disparity of the wedges (crossed and uncrossed; **Fig. 1b**). At a given time point, all wedges were presented the same disparity. In the centre of the wedge field, we presented a fixation square (side length = 1°) paired with horizontal and vertical nonius lines.

Two conditions were run: single- and mixed-polarity. In the single-polarity condition, the stimulus comprised uniform polarity dots, and alternated every presentation. In the mixed-polarity condition, the stimulus comprised equal proportions of randomly interspersed black and white dots (**Fig. 1b**). Read and Cumming (2018) have proposed that the mixed-polarity benefit arises from differences in the interocular cross correlation between some single- and mixed-polarity RDS. The difference in correlation arises from an interaction between the range of luminance in the stereograms and the variability of the binocular disparity. To avoid this potential confound, we designed the stimuli such that the variability of the binocular disparity in the images was low; binocular disparity was either zero or ±10 arcmin. **Figure 1c** shows the results of a comparison between image correlation for stimuli used in the single- and mixed-polarity conditions, confirming that there was no significant difference.

**Figure 1.**
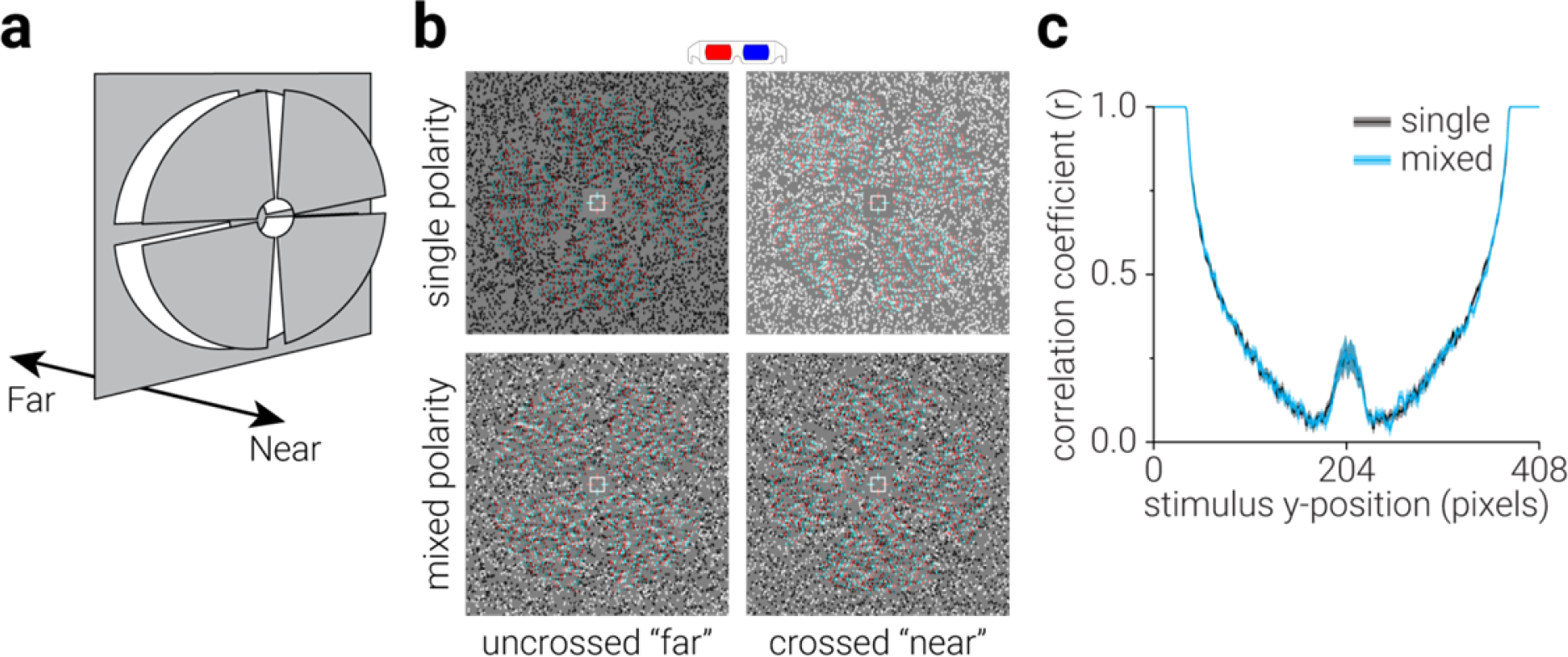
Stimuli used in the experiment. (**a**) Diagram of the depth arrangement of the stimuli; four disparity-defined wedges were simultaneously presented at either ±6 arcmin. (**b**) Examples of the near and far depth stimuli used in the single- and mixed-polarity conditions, designed for red-cyan anaglyph viewing. (**c**) The average left- and right-eye image cross-correlation as a function of stimulus y-axis position for single- and mixed-polarity RDS stimuli across a run (96 pairs of images). A paired t-test comparison between the cross-correlation across all positions was non-significant (single-polarity mean, .58; mixed-polarity mean, .58; *t*_95_=.82, *P*=.413). Shaded regions show s.e.m.

### Vernier task

During active (single/mixed-polarity condition) scans, participants performed an attentionally demanding Vernier task at fixation. This task served two purposes: it (i) ensured consistent attentional allocation between conditions, and (ii) provided a subjective measure of eye position, allowing us to assess whether there were any systematic differences in eye vergence between conditions. Participants were instructed to fixate a central cross hair fixation marker. The fixation marker consisted of a white square outline (side length 30 arcmin) and horizontal and vertical nonius lines (length 22 arcmin). One horizontal and one vertical line were presented to each eye in order to promote stable vergence and to provide a reference for a Vernier task (Popple, Smallman, & Findlay, 1998). The Vernier target line subtended 6.4 arcmin in height by 2.1 arcmin in width and was presented at seven evenly spaced horizontal offsets of between ±6.4 arcmin for 500 ms (with randomized onset relative to stimulus) on 33% of presentations. Participants were instructed to indicate, by button press, which side of the central upper vertical nonius line the target appeared, and the target was presented monocularly to the contralateral eye.

### Procedure

Participants underwent four MR spectroscopic acquisitions: an initial resting acquisition, followed by two active acquisitions, separated by a second half-length resting acquisition. The primary purpose of the second resting acquisition was to allow metabolite concentrations to return to a baseline state between active acquisitions. During resting acquisitions, participants were instructed to close their eyes. During active acquisitions, participants performed the Vernier task while viewing either single or mixed-polarity stereograms. The order of (single/mixed) active acquisitions was counterbalanced across participants.

### Magnetic resonance spectroscopy

Magnetic resonance scanning was conducted on a 3T Siemens Prisma equipped with a 32-channel head coil. Anatomical T1-weighted images were acquired for spectroscopic voxel placement with an ‘MP-RAGE’ sequence. For detection of GABA, spectra were acquired using a macromolecule-suppressed MEGA-PRESS sequence: TE=68ms, TR=3000ms; 256 transients of 2048 data points were acquired in 13 min experiment time; a 14.28 ms Gaussian editing pulse was applied at 1.9 (ON) and 7.5 (OFF) ppm; water unsuppressed 16 transients. Water suppression was achieved using variable power with optimized relaxation delays (VAPOR) and outer volume suppression (OVS). Automated shimming followed by manual shimming was conducted to achieve approximately 12 Hz water linewidth.

Spectra were acquired from a location targeting early visual cortex, i.e., V1/V2 (**Fig. 2a**). The voxel (30×30×20 mm) was placed medially in the occipital lobe, the lower face aligned with the cerebellar tentorium and positioned so to avoid including the sagittal sinus and to ensure it remained within the occipital lobe. To assess the consistency of voxel placement across subjects, the voxel location was used to mask a three-dimensional anatomical image of the observers’ brain. The resulting image was then transformed into Montreal Neurological Institute (MNI) space and the degree of overlap between voxel masks across was calculated (**Fig. 2b**). Further, we quantified the range of grey matter, white matter, and cerebral spinal fluid in each voxel (**Fig. 2c**).

Spectral quantification was conducted with GANNET (Baltimore, MD, USA), a MATLAB toolbox designed for analysis of GABA MEGA-PRESS spectra, modified to fit a double-Gaussian to GABA and Glx peaks. Individual spectra were frequency and phase corrected. Total creatine (tCR) signal intensity was determined by fitting a single mixed Gaussian-Lorentzian peak to the mean non-edited spectra, while water (H_2_O) signal intensity was determined by fitting a single mixed Gaussian-Lorentzian peak to the mean of the 16 water unsuppressed transients. ‘ON’ and ‘OFF’ spectra were subtracted to produce the edited spectrum, from which GABA and Glx signal intensity were modelled off double-Gaussian peaks. Intensities of GABA and Glx were normalized to the commonly used internal reference tCR (Jansen, Backes, Nicolay, & Kooi, 2006), yielding relative concentration values (i.e., GABA:Cr & Glx:Cr). The tCr signal is acquired within the same MEGA-PRESS acquisitions as the target metabolites; thus, normalization of GABA and Glx to tCr minimizes the influence of subject movement during the scan and eliminates the effects of chemical shift displacement (Mullins et al., 2014). Further, if metabolic changes during the acquisition occur globally, this will produce no change in the ratio of target metabolites to tCr. As an additional control, GABA and Glx were also normalized to H_2_O.

The fitting residual for tCr and GABA were divided by the amplitude of their fitted peaks to produce normalized measures of uncertainty. The quadratic of these was calculated to produce a combined measure of uncertainty for each measurement (Mullins et al., 2014; Rowland et al., 2016). This combined fitting residual was relatively low across all participants for all voxel locations; from 4.7% to 10.7% (mean: 7.8% ±0.1% s.e.m).

For the dynamic analysis, we used a sliding window (width, 128 acquisitions; step size, 1 acquisition) to measure average (**a**) GABA and (**b**) Glx concentration as it changed while participants viewed single- and mixed-polarity stimuli (256 acquisitions/13 minutes). Reducing the number of acquisitions in the averaged spectra reduced the signal-to-noise ratio. Thus, prior to running the dynamic analysis, metabolite concentration data were screened to remove noisy and/or spurious quantifications. In particular, we removed data points that were >4 standard deviations from the mean at each time point. This resulted in the removal of zero data points from the GABA dataset and 0.3% of data points from the Glx dataset. To remove spurious significant differences in the time course between conditions, a cluster correction was applied. Clusters were defined by the sum of their constituent (absolute) *t*-values and compared to a null hypothesis distribution of clusters produced by shuffling the condition labels (1000 permutations). Clusters below the 95^th^ percentile of the null hypothesis distribution were disregarded.

**Figure 2.**
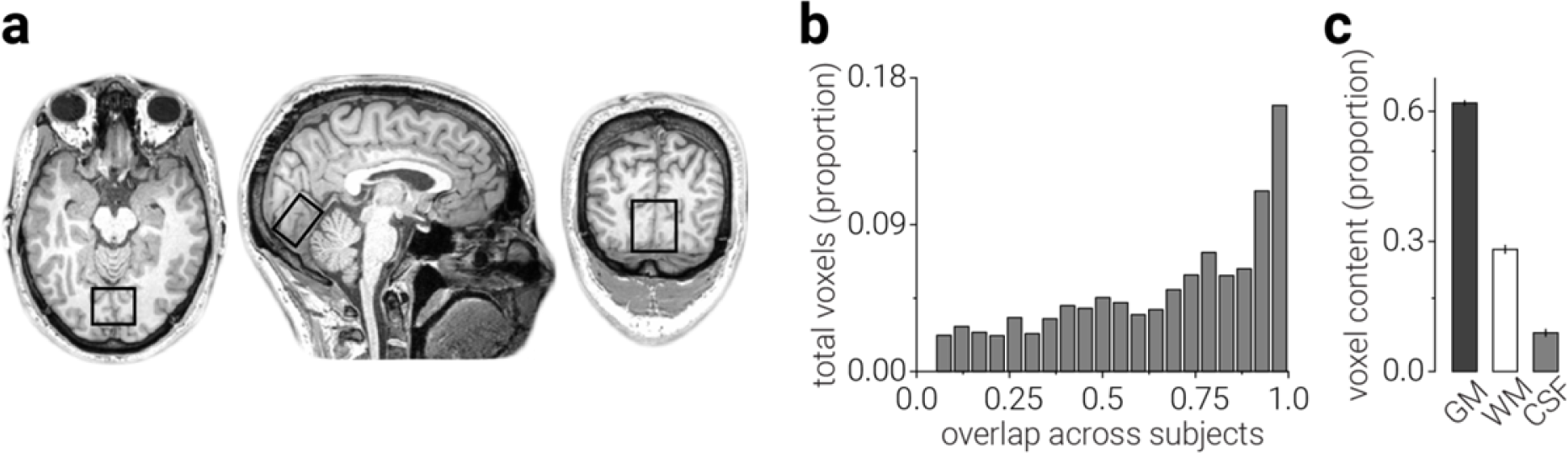
Magnetic resonance spectroscopy (MRS) voxel location across subjects. (**a**) A representative example of the location of the MRS voxel is shown in black. The location of the MRS voxel was highly consistent across subjects, as evidenced by (**b**) the overlap of voxels in MNI space and (**c**) the consistency of tissue content within the voxel. In (**c**), GM, WM, and CSF refer to grey matter, white matter, and cerebral spinal fluid, respectively.

## RESULTS

### GABA

Depth judgements of mixed-polarity stereograms appear to be more accurate than single-polarity stereograms, i.e., the mixed-polarity benefit (Harris & Parker, 1995). A recent study found that a deep neural network trained to classify depth in natural images also performed better on mixed-polarity than single-polarity stereograms (Goncalves & Welchman, 2017). Motivated by the observation that these stimuli evoked different patterns of positive and negative activity in the network, here we used magnetic resonance spectroscopy to test for potential changes in excitatory and inhibitory neurotransmitter concentration in human observers while they viewed single- and mixed-polarity stereograms.

Our region of interest was a voxel targeting early visual cortex (V1, V2) where the initial stages of binocular disparity processing take place. The GABA peaks were clearly visible within the spectra at approximately 3 ppm (**Fig. 3a**). We quantified the concentration of the primary inhibitory neurotransmitter γ-aminobutyric acid (GABA) and found that GABA concentration was significantly lower while viewing mixed-polarity stereograms compared to single-polarity stereograms (paired t-test, *t*_19_=2.99, *P*=.008; **Fig. 3b, c**). A possible concern might be that the observed change in GABA:Cr concentration relates to changes in total creatine (tCR), rather than GABA. However, we found the same result when GABA was referenced to water (paired t-test, *t*_19_=2.89, *P*=.009).

**Figure 3.**
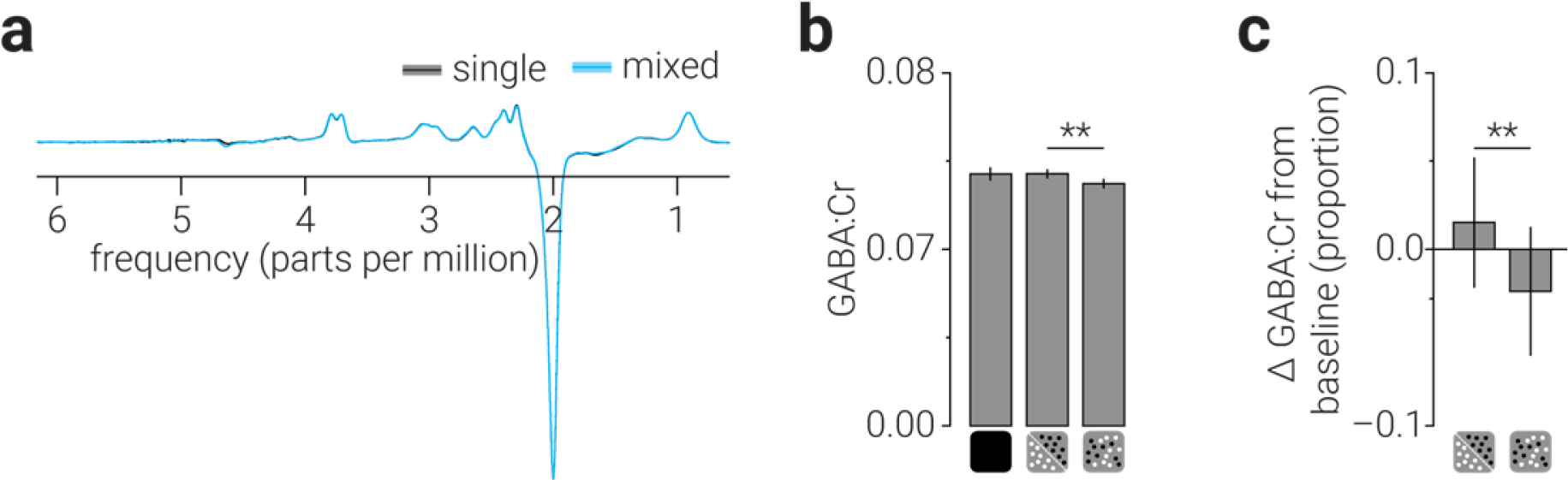
Average spectra and GABA measurements. (**a**) Spectra acquired while observes viewed single- and mixed-polarity random-dot stereograms (RDS), averaged across participants. (**b**) GABA concentration referenced to total creatine, from a voxel targeting early visual cortex, while observes were at rest or viewing single- and mixed-polarity RDS. (**c**) Same as (**b**), but referenced to GABA concentration at rest. Shaded regions in (**a**) and error bars in (**b**, **c**) indicate s.e.m. ** indicate significant differences with *P*<.01.

In order to assess the direction of metabolic change from “baseline” that viewing single- and mixed-polarity RDS produced, we acquired an initial resting measurement where observers were instructed to close their eyes. Despite the significant difference in GABA observed between viewing single-/mixed-polarity stereogram, the concentrations measured in these conditions did not significantly differ from the rest measurement (**Fig. 3b, c**).

An explanation for this might be that there was variability in the signal during the resting acquisition that was not present during the active acquisitions. For example, fixation and attention were controlled in the single- and mixed-polarity conditions by requiring observers to perform a demanding Vernier task at fixation. By contrast, we instructed observers to close their eyes during the resting acquisition, but we can neither confirm the extent to which they heeded this instruction, nor the focus of their attention. The variance in the resting condition was numerically larger in the resting conditions than the active conditions, but not significantly. However, we found that the correlation between GABA concentration while viewing single- and mixed-polarity RDS (Pearson correlation, *r*=.75, *P*=1.4e^−5^; **Fig. 4a**) was significantly higher than the that between resting GABA and active GABA (single: *z*=2.55, *P*=.01; mixed: *z*=2.56, *P*=.01), the latter of which were not significantly different from zero (single: *r*=.09, *P*=.70; mixed: *r*=.09, *P*=.70; **Fig. 4b, c**). That is, individuals’ GABA concentration was similar between “active” conditions, but it was not similar between active and rest conditions. This could be caused by additional sources of variability in the rest condition, and may explain why we did not detect a significant change in GABA concentration between rest and active conditions.

**Figure 4.**
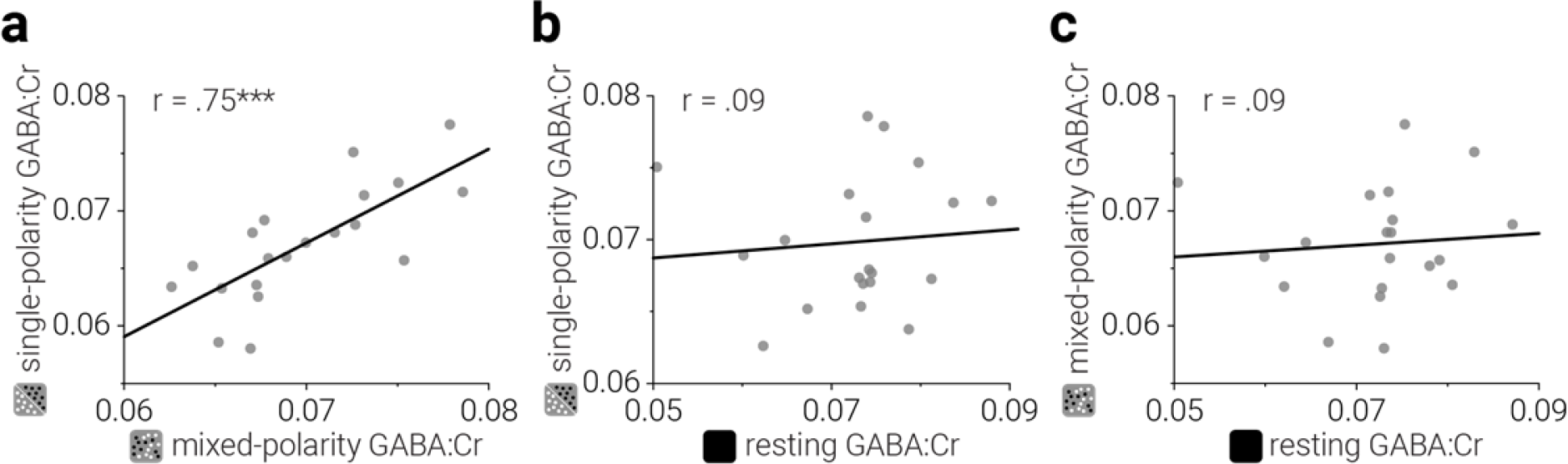
Within-subject correlations of GABA concentration in the early visual cortex while resting or viewing single/mixed-polarity random-dot stereograms. To assess within-subject consistency of metabolite concentrations, we compared GABA correlations between (**a**) active conditions, and (**b, c**) active conditions and at rest. *** indicate correlation scores with *P*<.001.

A possible concern might be that the observed difference in GABA concentration between active conditions was due to differences in attentional allocation or eye-movements while viewing single-/mixed-polarity stereograms. However, we found no evidence of a difference in performance on the attentionally demanding Vernier task between conditions, either in accuracy (paired t-test, *t*_18_=0.93, *P*=.36; **Fig. 5a**) or response time (paired t-test, *t*_18_=0.60, *P*=.57; **Fig. 5b**). For each participant we fitted a cumulative Gaussian to the proportion of “right” responses as a function of the horizontal offsets of the targets, to obtain bias measurements. Bias (deviation from the desired vergence position) in participants’ judgments was not significantly different between conditions (paired *t*-test, *t*_18_=0.31, *P*=.76; **Fig. 5c**), or from zero (paired *t*-test, single: *t*_18_=1.34, *P*=.20; mixed: *t*_18_=1.36, *P*=.19). Performance on the Vernier task therefore suggests that participants were able to maintain stable eye vergence equally well between single- and mixed-polarity conditions (Popple et al., 1998).

**Figure 5.**
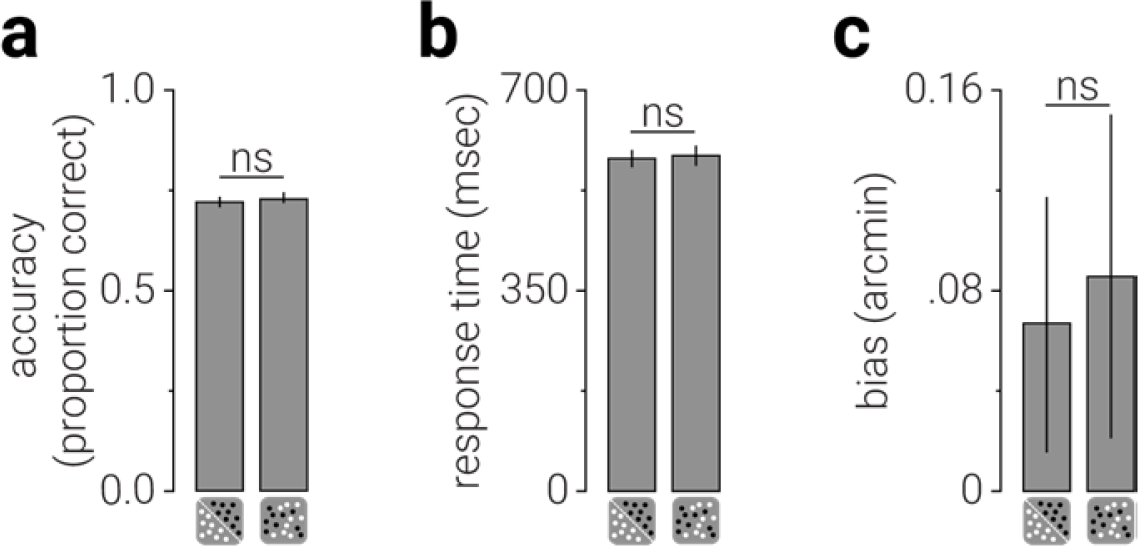
Vernier task performance during single- and mixed-polarity viewing acquisitions. Comparison between (**a**) accuracy and (**b**) response time on a Vernier task performed while observers viewed single- and mixed-polarity stimuli. (**c**) Bias (deviation from the desired vergence position) in participants’ judgments. Error bars indicate s.e.m.

Having established that viewing single- and mixed-polarity stereograms produced differences in GABA concentration in the early visual cortex, we tested whether the difference in GABA between conditions was stable or changed during the presentation. We used a sliding window to measure GABA:Cr as it dynamically changed over the course of the acquisition (Branzoli, Techawiboonwong, Kan, Webb, & Ronen, 2013; Schaller, Mekle, Xin, Kunz, & Gruetter, 2013). We found that the difference in concentration while viewing single-/mixed-polarity stereograms was greatest early in the acquisition (*P*<.05 from step 2-22, *P*_min_=5.4e^−4^; **Fig. 6**). This suggests that visual stimulation altered GABA concentration most during the first half of the presentation (~7 min), after which it began to return to its initial state.

**Figure 6.**
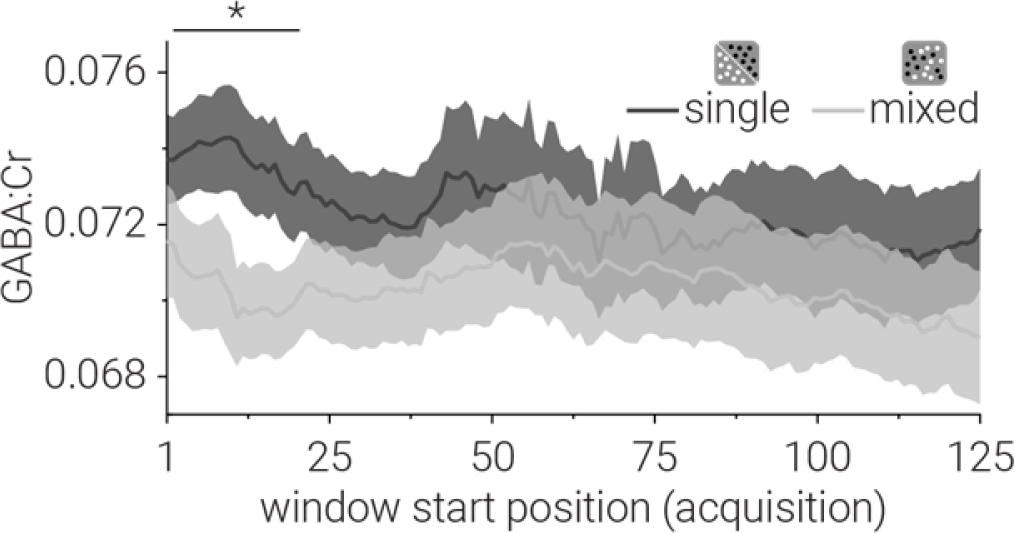
Dynamic GABA concentration in early visual cortex. We used a sliding window (kernel size, 128 acquisitions; step size, 1 acquisition) to measure GABA concentration as it changed while participants viewed single- and mixed-polarity stimuli (256 acquisitions/13 minutes). The horizontal line at the top of the plot indicates periods where the difference in concentration while viewing single-and mixed-polarity stimuli was significant, following cluster correction. Shaded regions indicate s.e.m. * indicate continuous period of significant differences with *P*<.05.

### Glx

We found that GABA concentration measured from a voxel targeting V1 and V2 was different when observers viewed single- and mixed-polarity stereograms, suggesting that there was different involvement of inhibitory systems in the processing of these stimuli. We then compared the concentration of Glx, a complex comprising Glutamate (the primary excitatory neurotransmitter) and Glutamine, between these conditions. The Glx peaks were clearly visible in the spectra at approximately 3.8 ppm (**Fig. 3a**). We found that Glx was significantly higher in the mixed-polarity condition (paired t-test, *t*_19_=2.35, *P*=.029; **Fig. 7a, b**). Further, we found the same result when Glx was referenced to water (paired t-test, *t*_19_=2.47, *P*=.023).

**Figure 7.**
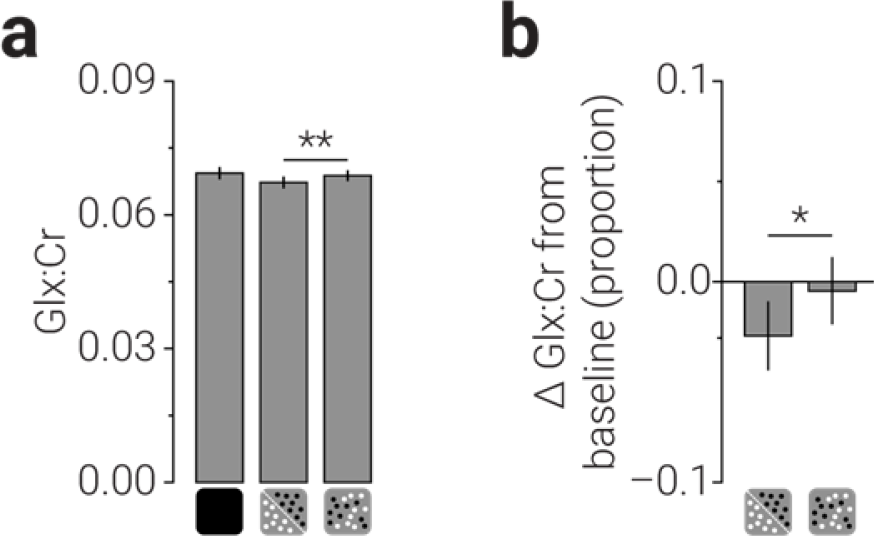
Glx measurements. (**a**) Glx concentration referenced to total creatine, from a voxel targeting early visual cortex, while observes were at rest or viewing single- and mixed-polarity RDS. (**b**) Same as (**a**), but referenced to Glx concentration at rest. Error bars indicate s.e.m. ** indicate significant differences with *P*<.01.

Similar to the GABA results, comparison of Glx measured in the viewing conditions to that measured at rest yielded no significant differences. However, unlike the GABA results, the correlation between Glx concentration while viewing single- and mixed-polarity RDS (Pearson correlation, *r*=.86, *P*=9.0e^−7^; **Fig. 8a**) was not significantly higher than the that between resting Glx and active Glx (single: *z*=1.79, *P*=.073; mixed: *z*=1.76, *P*=.078; **Fig. 8b, c**). This may be because Glutamate is only a component of the Glx complex, and the other component (Glutamine) may be more stable between resting and active periods. For the dynamic analysis of Glx concentration, we found that the difference remained stable during the acquisition (**Fig. 9**).

**Figure 8.**
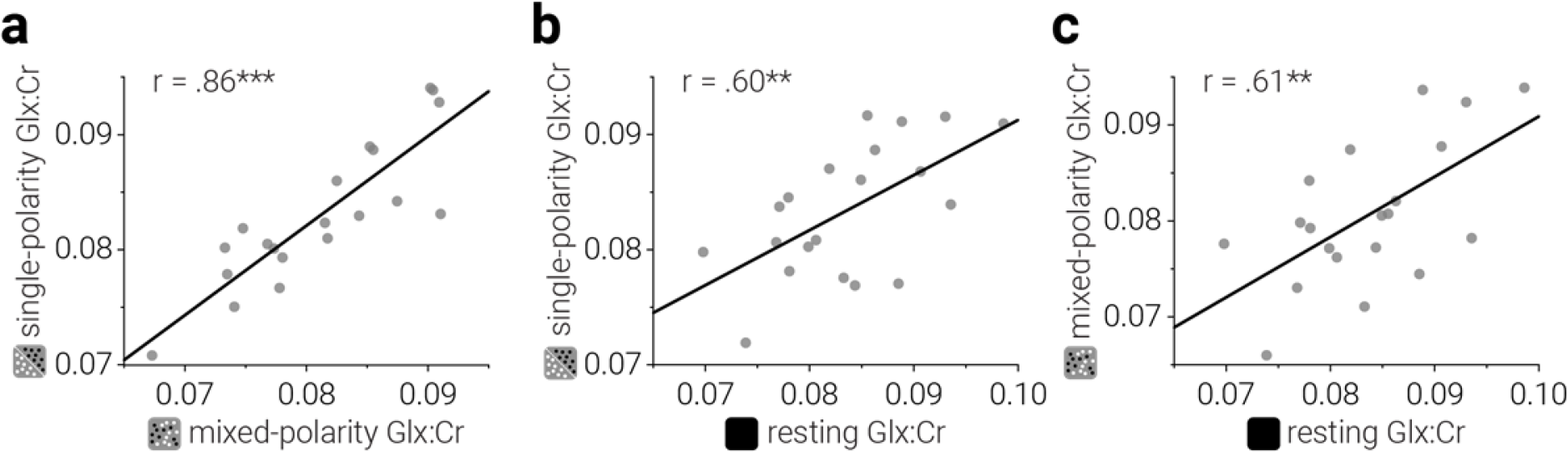
Within-subject correlations of Glx concentration in the early visual cortex while resting or viewing single/mixed-polarity random-dot stereograms. To assess within-subject consistency of metabolite concentrations, we compared Glx correlations between (**a**) active conditions, and (**b, c**) active conditions and at rest.

**Figure 9.**
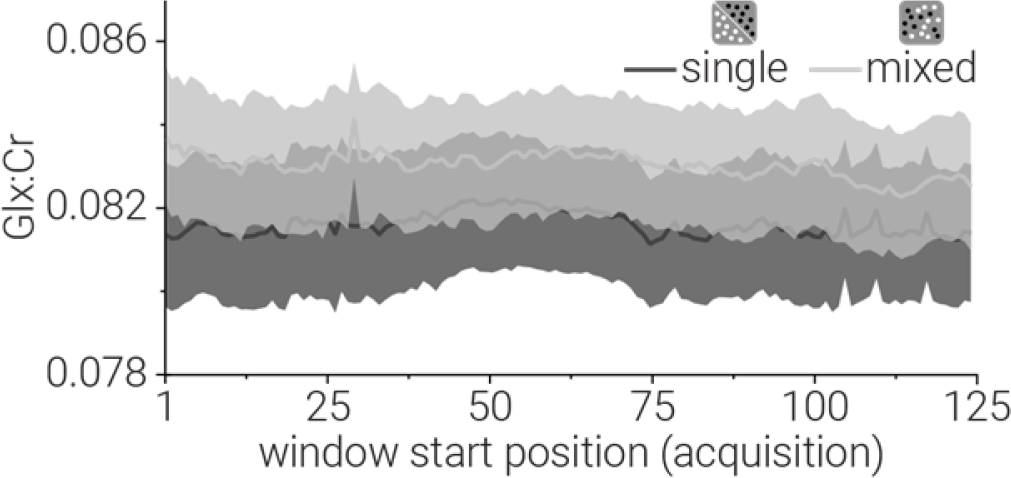
Dynamic Glx concentration in early visual cortex. We used a sliding window (kernel size, 128 acquisitions; step size, 1 acquisition) to measure GABA concentration as it changed while participants viewed single- and mixed-polarity stimuli (256 acquisitions/13 minutes). No significant differences survived cluster correction. Shaded regions indicate s.e.m.

We found that concentrations of GABA in the early visual cortex were significantly lower when subjects viewed mixed-polarity stereograms, compared to when viewing single-polarity stereograms. By contrast, we found the opposite effect for Glx; concentrations were higher when viewing mixed-polarity stereograms. It is possible that a decrease in one may have coincided with an increase in the other, e.g., to maintain a particular balance of inhibition and excitation. However, we found no evidence for a correlation between the difference in GABA and Glx between single- and mixed-polarity viewing conditions (Pearson correlation, *r*=.−14, *P*=.56; **Fig. 10**).

**Figure 10.**
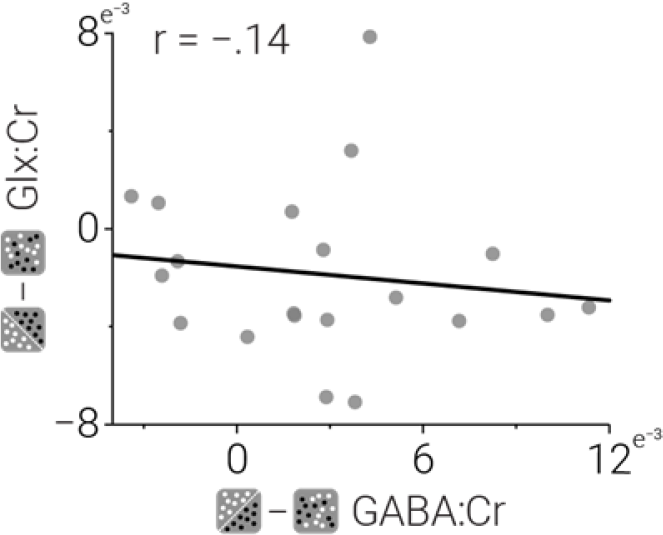
The difference in Glx:Cr between single- and mixed-polarity viewing conditions as a function of the difference in GABA:Cr between single- and mixed-polarity viewing conditions.

## DISCUSSION

Depth judgements are more accurate when binocular images are composed of both light and dark features, rather than just one or the other (Harris & Parker, 1995; Read, Vaz, & Serrano-Pedraza, 2011). This finding was initially interpreted as evidence of separate ON and OFF binocular channels; however, contradictory physiological (Schiller, 1992), behavioural (Read et al., 2011) and theoretical (Goncalves & Welchman, 2017; Read & Cumming, 2018) evidence has cast doubt on this hypothesis. Here we tested the potential for changes in the balance of excitation and inhibition that are produced by viewing these single- and mixed-polarity random-dot stereograms (RDS). This follows directly from the recent observation that these stimuli evoke different levels of positive and negative activity in a deep neural network, which also showed the mixed-polarity benefit (Goncalves & Welchman, 2017). We show that GABA concentration measured in the early visual cortex is lower when viewing mixed-polarity stereograms than single-polarity stereograms. Further, we show the opposite pattern of results for the Glutamate/Glutamine complex (Glx).

The finding that GABA concentration was different when subjects viewed single- and mixed-polarity stereograms indicates that these stimuli evoke different levels of suppressive activity in the early visual cortex. Given the current understanding of MRS-measured changes in GABA and its relationship to neural function, interpreting these results as unambiguous evidence for increased suppressive activity in one condition or the other is challenging. For example, increased suppressive activity demands a corresponding increase in GABA synthesis, which may expand the “pool” of GABA that can be detected by MRS. However, when GABA is released from the cell body into the synapse it becomes bound to GABA receptors, which broadens its resonance and makes it less detectable with MRS (Jahnke et al., 2002; Xu, Seto, Tang, & Firestone, 2000). While the former point predicts an increase in MRS-measured GABA concentration, the latter predicts a reduction. Establishing the directionality of the GABA change is further complicated as we did not detect a metabolic difference in either viewing condition from baseline.

In addition to a difference in GABA concentration, we also observed a difference in Glx (a complex comprising Glu and Gln) between single-/mixed-polarity stereogram viewing conditions. Given that Gln is a primary source of GABA synthesis (Patel, Rothman, Cline, & Behar, 2001; Paulsen, Odden, & Fonnum, 1988; Rae et al., 2003), the change we observed in Glx may be due to differences in the activity of the inhibitory system, as evidenced by the change in GABA concentration. However, we found no evidence supporting this interpretation; that is, there was no relationship between the magnitude of change in these metabolites between subjects. Alternatively the difference in Glx may be attributed to altered metabolic and/or neurotransmitter synthesis of Glu due to differences in activity of the excitatory system while viewing single-/mixed-polarity stereograms. This explanation is supported by MRS work with phantoms that estimate the signal contribution to Glx in the MEGA-PRESS difference spectrum as either equal parts Glu and Gln (van Veenendaal et al., 2018) or primarily Glu (Shungu et al., 2013).

These results are broadly consistent with the recent observation that single- and mixed-polarity stereograms evoke different levels of positive and negative activity from a convolutional neural network trained on natural stereo images to make depth judgements (Goncalves & Welchman, 2017). Critically, this neural network also reproduced the mixed-polarity benefit, suggesting that the difference in positive/negative activity may underlie this puzzling phenomenon.

Read and Cumming (2018) have proposed that the mixed-polarity benefit arises from subtle changes in image correlation under some restricted circumstances. Given that the image correlation was the same between single- and mixed-polarity conditions, the difference in neurotransmitter concentration cannot be attributed to this hypothesis. It is also unlikely that differences in attention between conditions provide an explanation for the changes in metabolic concentrations because we found equivalent performance for the psychophysical task that participants performed at the fixation marker. We cannot, however, rule out the possibility that the observed effects were caused by monocular differences between the stimuli. For instance, the luminance of the stimuli in the single- and mixed-polarity conditions was the same when averaged over any two consecutive presentations, i.e., all white followed by all black is equivalent to two mixed presentations, but between consecutive presentations the mean luminance of stimuli in the single-polarity condition varied more than in the mixed-polarity condition.

A limitation of MRS is that it measures total concentration of neurochemicals within a localized region and cannot distinguish between intracellular and extracellular pools of GABA. This is relevant, because these pools are thought to have different roles in neuronal function. Here we show that GABA concentration in the early visual cortex is different when viewing single- or mixed-polarity RDS, indicating changes in the level of intracellular vesicular GABA, which drives neurotransmission (Belelli et al., 2009). However, MRS also measures extracellular GABA, which maintains tonic cortical inhibition (Martin & Rimvall, 1993) and is unlikely to be altered by our experimental manipulations. While the magnitude of GABA difference we observed was consistent with previous work (Bednařík et al., 2015; Mekle et al., 2017), this may explain its meagre size (4%). The difference in Glx (2%) is also consistent with previous work (Bednařík et al., 2015; Mangia et al., 2007; Mullins, Rowland, Jung, & Sibbitt, 2005; Schaller et al., 2013).

Dynamic analysis of GABA and Glx revealed that the difference in GABA was largest early in the scan, whereas for Glx difference appeared to be relatively stable throughout. These results are broadly consistent with previous work (Bednařík et al., 2015; Chen et al., 2017; Lin, Stephenson, Xin, Napolitano, & Morris, 2012); however, whereas we found early differences in GABA, i.e., in the first 7 min of stimulation, (Chen et al., 2017) found the maximum difference after 5 min of hand-clenching. The distinct time courses found for GABA and Glx when viewing single-/mixed-polarity stereograms may reflect differences in the extent to which inhibitory/excitatory systems were engaged. In particular, the early difference in GABA may suggest a large difference in evoked inhibitory activity, which was subsequently accommodated. By contrast, the small but consistent difference in Glx may reflect a smaller, more stable, difference in evoked excitatory activity.

To summarize, here we find differences in GABA and Glx concentration when subjects view single- and mixed-polarity RDS. These results indicate different levels of inhibitory and excitatory activity are evoked by the stereoscopic computation of these stimuli and may hold the key to understanding why depth judgements are better for stereograms comprised of both light and dark features.

## Acknowledgements

This work was supported by the Leverhulme Trust (ECF-2017-573) to RR and the Wellcome Trust (095183/Z/10/Z) to AEW.

